# Spatial transcriptomics reveals D2-associated synaptic transcriptional attenuation in the chronically stressed dorsal striatum

**DOI:** 10.64898/2026.07.12.737112

**Authors:** Jinhee Bae, Heh-In Im

## Abstract

Chronic stress alters striatal functions involved in motivation, action selection, and behavioral adaptation, yet cell-type-associated transcriptional organization in the dorsal striatum remains unclear. We used RNAscope-guided GeoMx spatial transcriptomics to compare D1 and D2 neuronal compartments in matched dorsal striatal regions after chronic restraint stress (CRS). CRS engaged both populations and produced comparable numbers of differentially expressed genes. Gene set enrichment analysis revealed partially overlapping CRS-associated pathway attenuation in D1 and D2 neurons, indicating stress-responsive transcriptional organization in both populations. However, D2 responses showed more coherent convergence around receptor-trafficking and synaptic signaling programs, including AMPA receptor trafficking and EPHB-mediated signaling. Moreover, under the same threshold-defined DEG criteria, CRS-downregulated D2 genes resolved into synapse-centered functional annotation categories, including glutamatergic synapse, postsynaptic organization, and dendritic spine, whereas D1 gene sets did not show a comparable pattern. These findings provide a framework for comparing stress-associated D1/D2 transcriptional organization in the dorsal striatum.

**Graphical abstract:** 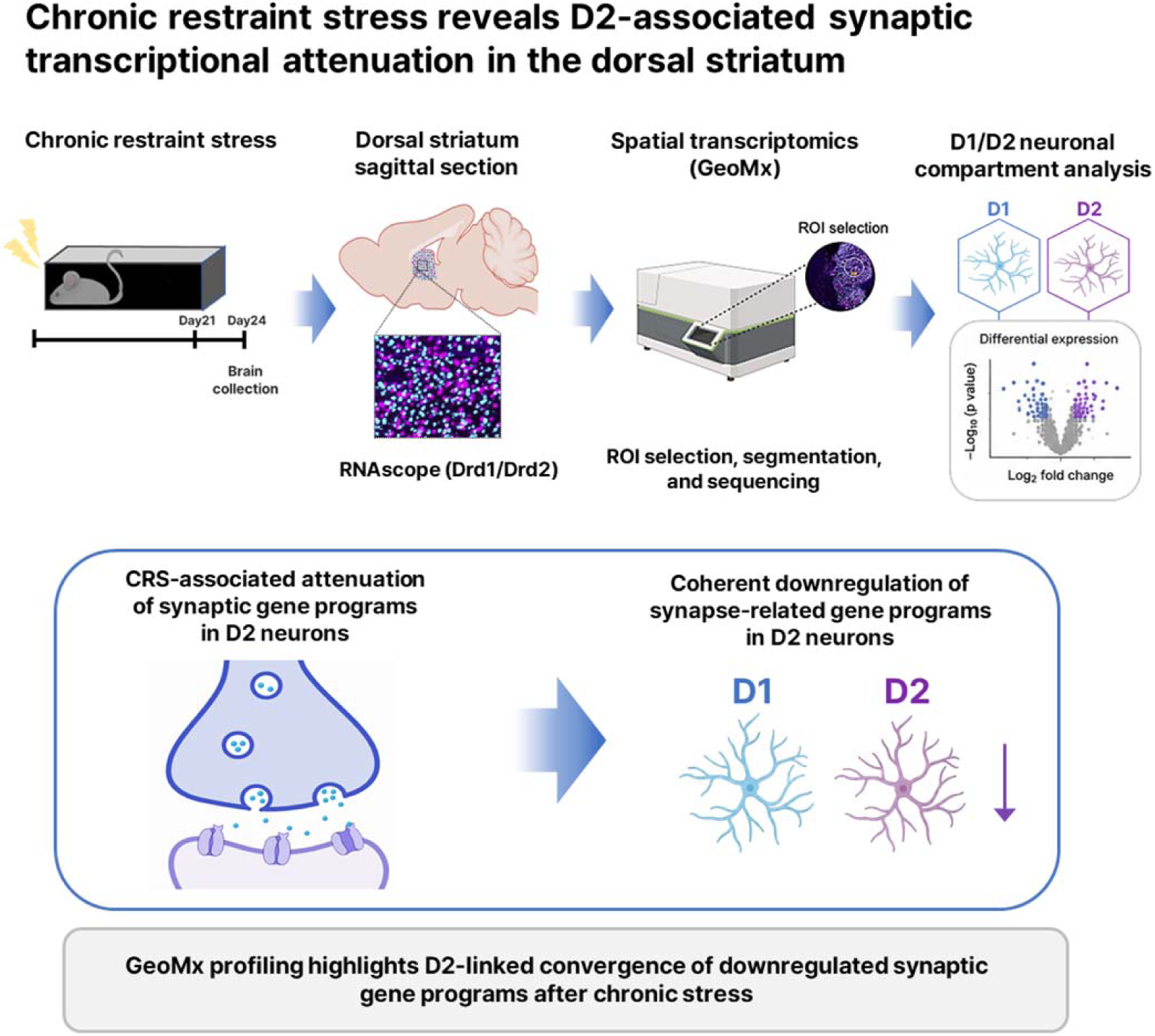

## Introduction

Stress-related depressive states involve disruptions in motivational, adaptive, and action-selection processes, functions closely linked to striatal systems^1–3^. Although the dorsal striatum has traditionally been studied for its roles in action selection, habit formation, motor control, and basal ganglia circuit function^4^, accumulating evidence from human imaging and experimental stress studies implicates this region in stress-related affective dysfunction and behavioral adaptation^5–8^. Recent advances in spatial transcriptomics have enabled transcriptome-wide profiling while preserving tissue organization, and recent applications have mapped striatal neuronal heterogeneity under baseline conditions^9,10^. However, stress-associated D1/D2 transcriptional organization within the dorsal striatum remains unresolved.

Human and animal studies support a role for the dorsal striatum in stress- and depression-related behavioral regulation. In patients with depression, altered dorsal corticostriatal and frontostriatal connectivity has been reported, and striatal dopamine deficits and impaired reward prediction-error encoding have been linked to altered striatal functional connectivity^1–3,5^. Experimental studies further suggest that dorsal striatal mechanisms contribute to stress coping, reward learning, and behavioral adaptation, including D2 receptor-associated coping responses, dorsal striatal prediction-error signaling, Shati/Nat8l–BDNF-related stress vulnerability, and chronic stress-induced striatal disinhibition^6–8,11^.

Despite these advances, how chronic stress reorganizes D1 and D2 transcriptional programs within the dorsal striatum remains poorly resolved. D1 and D2 neurons are spatially intermingled within this region but represent functionally distinct output populations^4,12^. Although previous molecular studies have provided important insights into striatal cell-type diversity^9,10^, stress-associated D1/D2 transcriptional organization has not been examined within matched dorsal striatal tissue regions. A spatially guided within-region strategy is therefore well suited to assess how chronic stress engages D1 and D2 neuronal populations in the dorsal striatum.

Here, we used RNAscope-guided GeoMx spatial transcriptomics to examine chronic stress-associated D1 and D2 neuronal transcriptional organization within matched dorsal striatal tissue regions. This within-region strategy allowed us to compare D1 and D2 neuronal populations in the same anatomical context and assess whether chronic stress engages these populations through shared or distinct transcriptional programs. We found that chronic restraint stress engaged both D1 and D2 neurons, with comparable numbers of differentially expressed genes across the two populations. Although CRS-associated pathway responses partially overlapped between D1 and D2 neurons, D2 responses showed more coherent organization around receptor-trafficking and synaptic signaling programs, including AMPA receptor trafficking and EPHB-mediated signaling. Together, these findings provide a spatially resolved framework for understanding how chronic stress reorganizes dorsal striatal D1 and D2 neuronal transcriptional states.

## Results

### RNAscope-guided GeoMx profiling resolves D1 and D2 neuronal compartments in the dorsal striatum

To examine cell-type-resolved transcriptional responses to chronic stress in the dorsal striatum, mice were exposed to chronic restraint stress (CRS) for 6 h per day over 21 days, and brains were collected 3 days after the final restraint session (Figure 1A). Dorsal striatal regions of interest were selected for spatial transcriptomic profiling based on anatomical landmarks (Figure 1B). We combined RNAscope detection of *Drd1* and *Drd2* transcripts with GeoMx spatial profiling to segment D1- and D2-enriched neuronal compartments within matched dorsal striatal tissue regions (Figure 1C). Representative RNAscope images showed clearly distinguishable *Drd1* and *Drd2* signals within the dorsal striatum, supporting the suitability of this strategy for D1/D2-informed segmentation (Figure S1).

**Figure 1.**
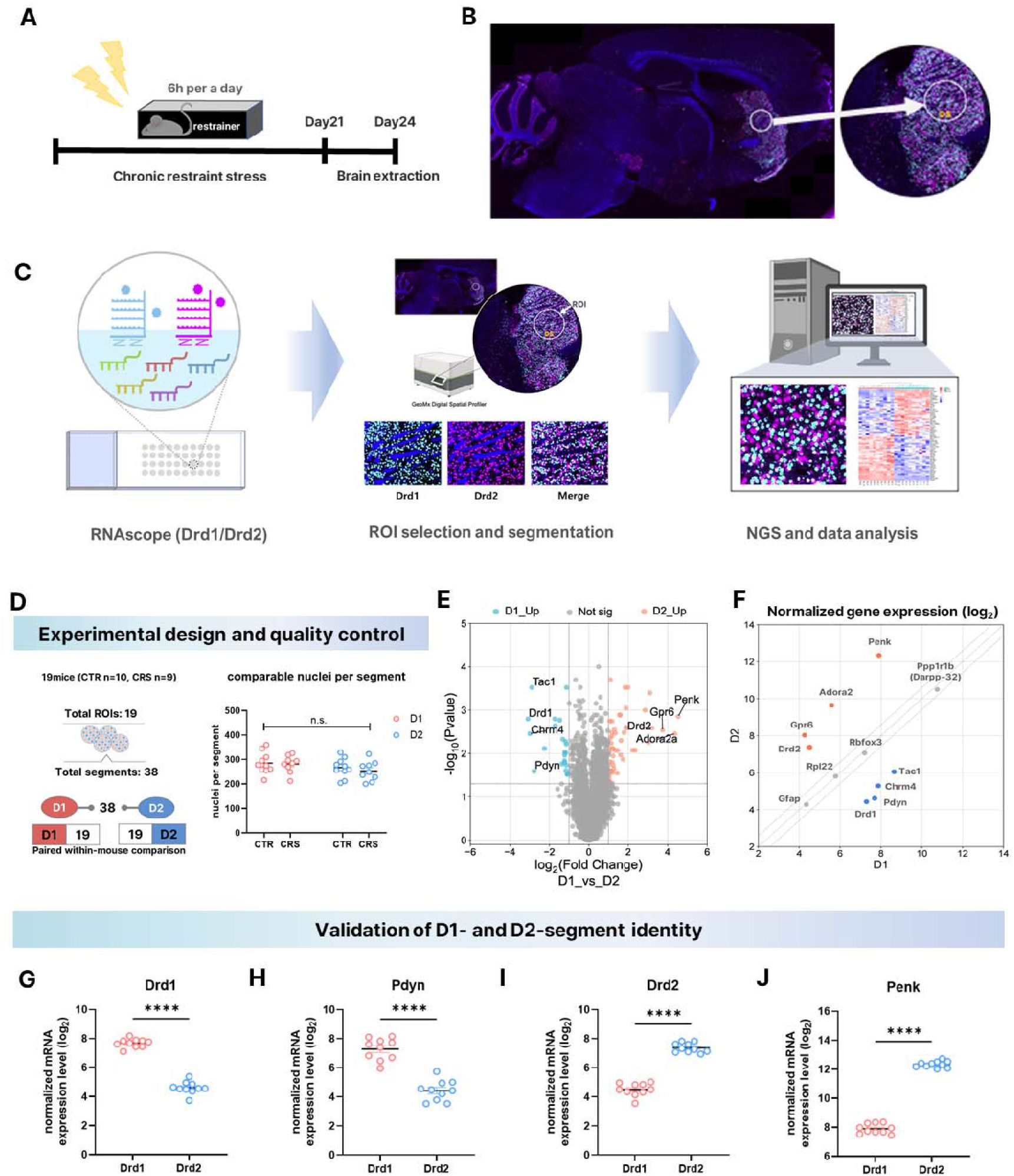
Experimental design and validation of D1/D2-segmented spatial transcriptomic profiling in the dorsal striatum following chronic restraint stress. (A) Schematic of the chronic restraint stress (CRS) paradigm. Mice were subjected to restraint stress for 6 h per day for 21 consecutive days, and brains were collected 3 days after the final restraint session. (B) Representative sagittal brain section showing the dorsal striatum (DS) region used for spatial transcriptomic analysis. The magnified image indicates the selected region of interest (ROI), with Drd1 signal shown in cyan, Drd2 signal in magenta, and nuclear staining in. (C) Overview of the RNAscope-guided GeoMx spatial transcriptomic workflow. RNAscope probes targeting *Drd1* and *Drd2* were used to label D1- and D2-enriched neuronal compartments, followed by ROI selection, D1/D2 molecular segmentation, next-generation sequencing (NGS), and downstream analysis. Representative single-channel and merged images show Drd1 in cyan, Drd2 in magenta, and nuclear staining in blue. (D) Experimental design and quality control. A total of 19 mice were analyzed, including 10 control and 9 CRS mice. Each mouse contributed one dorsal striatal ROI, which was segmented into paired D1 and D2 compartments, yielding 38 total segments. Nuclei counts did not differ significantly between control and CRS groups or between D1 and D2 segments, supporting comparable segment cellularity across groups. Statistical analysis was performed using a mixed-effects model with stress condition and segment identity (D1 vs. D2) as factors: stress effect, *F*(1, 17) = 0.4226, *p* = 0.5243; segment effect, *F*(1, 17) = 3.782, *p* = 0.0686; interaction, *F*(1, 17) = 0.1597, *p* = 0.6944. n.s., not significant. (E) Volcano plot showing differential gene expression between D1- and D2-enriched segments across all samples. Representative D1- and D2-associated marker genes are labeled. Colored points indicate genes enriched in D1 or D2 segments, and gray points indicate genes that were not significantly different between D1 and D2 segments. (F) Scatter plot comparing normalized gene expression levels between D1 and D2 segments. Genes near the diagonal line represent transcripts with comparable expression between D1 and D2 segments, including broadly expressed and glial-associated genes, whereas canonical D1- and D2-enriched markers deviate toward their expected compartments. (G–J) Validation of D1- and D2-segment identity using canonical marker genes in paired D1 and D2 segments from control mice. D1-associated genes *Drd1* and *Pdyn* were enriched in D1 segments, whereas D2-associated genes *Drd2* and *Penk* were enriched in D2 segments. Data are shown as log□-normalized expression values and presented as mean ± SEM. Statistical significance was determined by paired two-tailed Student’s *t* test: *Drd1*, *t*(9) = 20.70, *p* < 0.0001; *Pdyn*, *t*(9) = 20.94, *p* < 0.0001; *Drd2*, *t*(9) = 20.97, *p* < 0.0001; *Penk*, *t*(9) = 52.41, *p* < 0.0001. \*\*\*\**p <* 0.0001.

The dataset included 19 mice, comprising 10 control and 9 CRS mice. Each mouse contributed one dorsal striatal region of interest (ROI), which was segmented into paired D1 and D2 compartments, yielding 38 total segments. Nuclei counts did not differ significantly between control and CRS groups or between D1 and D2 segments, supporting consistent sampling and comparable segment cellularity across experimental groups (Figure 1D).

We next validated the molecular identity of the segmented compartments. Differential expression analysis between D1- and D2-enriched segments showed the expected distribution of canonical striatal markers, with D1-associated genes such as *Drd1*, *Pdyn*, *Tac1*, and *Chrm4* enriched in D1 segments, and D2-associated genes such as *Drd2*, *Penk*, *Gpr6*, and *Adora2a* enriched in D2 segments (Figure 1E and Table S1). Normalized gene expression profiles further showed clear enrichment of canonical D1 and D2 markers in their corresponding segmented populations (Figure 1F). Additional marker-based analysis under baseline conditions confirmed subtype-specific molecular signatures of D1 and D2 segments (Figure S2). Direct comparison of representative markers confirmed robust enrichment of *Drd1* and *Pdyn* in D1 segments and *Drd2* and *Penk* in D2 segments (Figures 1G–1J). Baseline functional annotation and network analyses under control conditions further supported molecular distinctions between D1 and D2 segments, while showing that shared functional labels could be supported by distinct gene compositions (Figures S3 and S4; Tables S2 and S3).

Together, these results validate the RNAscope-guided D1/D2 segmentation strategy and establish a reliable spatial transcriptomic dataset for comparing CRS-associated transcriptional responses between dorsal striatal D1 and D2 neuronal populations.

### CRS induces comparable DEG numbers but partially distinct transcriptional responses in D1 and D2 neurons

Having validated RNAscope-guided D1/D2 segmentation, we next compared CRS-associated transcriptional responses in D1 and D2 neurons. Threshold-based differential expression analysis was performed using genes with at least a 1.3-fold change and nominal *p* < 0.05, corresponding to |log□FC| ≥ 0.3785. This analysis identified similar numbers of CRS-responsive genes in the two neuronal populations, with 369 DEGs in D1 neurons and 357 DEGs in D2 neurons (Figure 2A; Tables S4 and S5). In both populations, most CRS-responsive genes were upregulated, whereas smaller subsets were downregulated: 330 upregulated and 39 downregulated genes in D1 neurons, and 313 upregulated and 44 downregulated genes in D2 neurons. These results indicate that CRS induced comparable numbers of DEGs in D1 and D2 neurons, suggesting that cell-type-related differences were not simply explained by the overall number of CRS-responsive genes.

**Figure 2.**
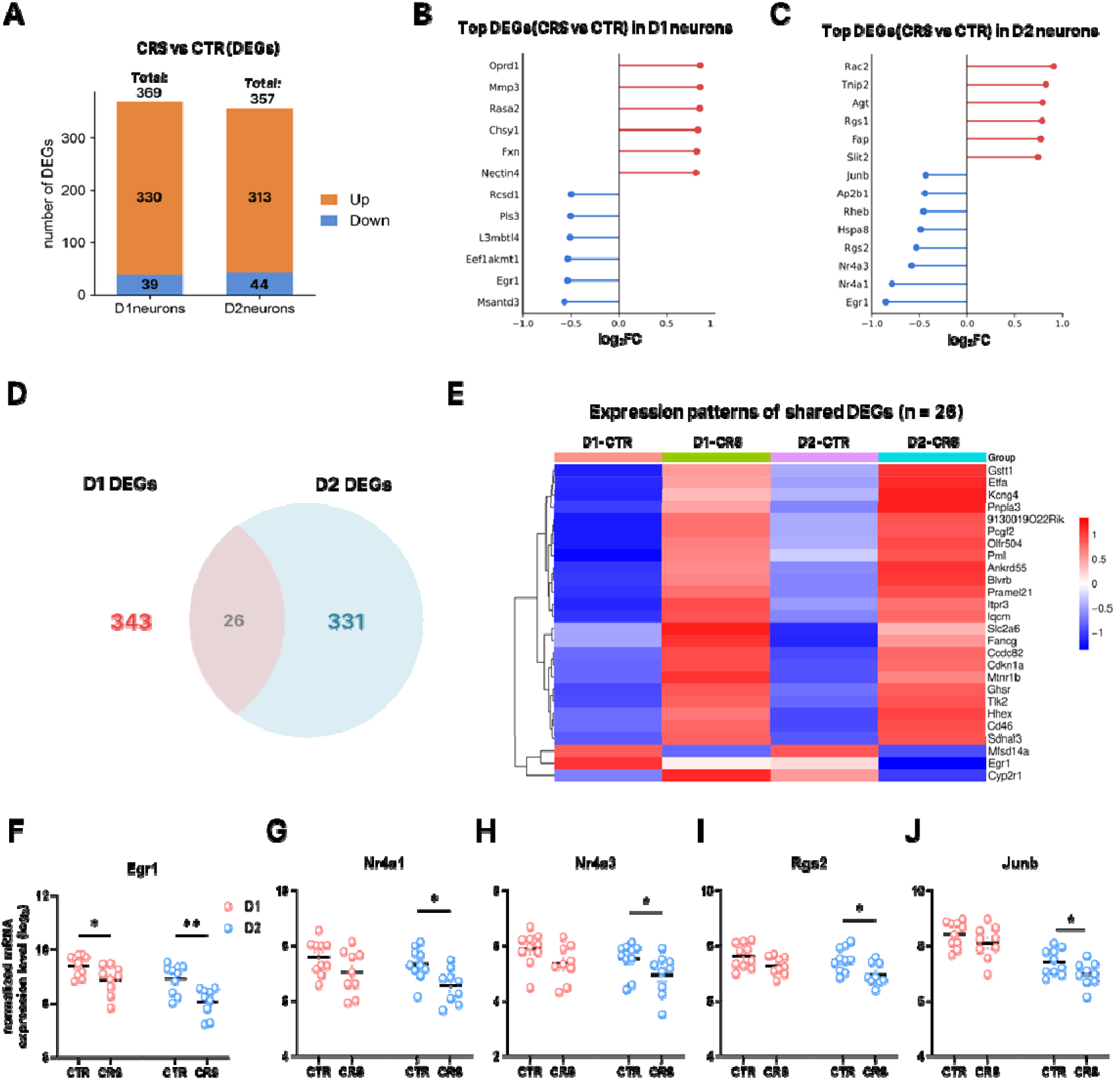
CRS induces comparable DEG numbers but partially distinct transcriptional responses in D1 and D2 neurons. (A) Summary of CRS-responsive differentially expressed genes (DEGs) identified in D1 and D2 neuronal segments. DEGs were defined using threshold-based criteria of |log□FC| ≥ 0.3785, corresponding to at least a 1.3-fold change, and nominal *p* < 0.05. D1 neurons showed 330 upregulated and 39 downregulated genes, whereas D2 neurons showed 313 upregulated and 44 downregulated genes. (B and C) Lollipop plots showing selected representative CRS-responsive genes from the top-ranked upregulated and downregulated DEG lists in D1 neurons (B) and D2 neurons (C). Genes were selected to illustrate major CRS-associated expression patterns in each neuronal population. Red indicates upregulated genes and blue indicates downregulated genes in CRS relative to control. (D) Venn diagram showing the overlap of CRS-responsive DEGs between D1 and D2 neurons. A total of 26 DEGs were shared between the two neuronal populations, whereas 343 were D1-only and 331 were D2-only DEGs. (E) Heatmap showing expression patterns of the 26 shared CRS-responsive DEGs across D1 and D2 neuronal segments under control and CRS conditions. Rows represent genes, and columns represent group-averaged expression values for D1-control, D1-CRS, D2-control, and D2-CRS. Color indicates row-scaled normalized expression. (F–J) Individual gene-level visualization of representative activity-related genes identified among CRS-downregulated genes. *Egr1* showed CRS-associated reduction in both D1 and D2 neurons (F), whereas *Nr4a1* (G), *Nr4a3* (H), *Rgs2* (I), and *Junb* (J) showed more evident CRS-associated decreases in D2 neurons. Data are shown as log□-normalized expression values and presented as mean ± SEM. Statistical analysis was performed using a mixed-effects model with stress condition and neuronal population as factors, followed by post hoc comparisons as indicated in the graphs. Full mixed-effects model statistics and post hoc comparisons are provided in Tables S6 and S7. \**p <* 0.05, \*\**p <* 0.01.

We then visualized selected representative CRS-responsive genes from the top-ranked DEG lists to examine major expression patterns in each neuronal population. In D1 neurons, representative upregulated genes included *Oprd1*, *Mmp3*, *Rasa2*, *Chsy1*, *Fxn*, and *Nectin4*, whereas downregulated genes included *Rcsd1*, *Pls3*, *L3mbtl4*, *Eef1akmt1*, *Egr1*, and *Msantd3* (Figure 2B and Table S4). In D2 neurons, representative upregulated genes included *Rac2*, *Tnip2*, *Agt*, *Rgs1*, *Fap*, and *Slit2*, whereas several downregulated genes were associated with activity-related transcriptional and signaling responses, including *Junb*, *Rgs2*, *Nr4a3*, *Nr4a1*, and *Egr1* (Figure 2C and Table S5). These representative gene-level patterns suggested that CRS responses contained both shared and cell-type-associated components.

Consistent with this observation, overlap analysis showed that only a subset of CRS-responsive genes was shared between D1 and D2 neurons. Among the identified DEGs, 26 genes overlapped between the two neuronal populations, whereas 343 were D1-only and 331 were D2-only DEGs (Figure 2D; Tables S4 and S5). Heatmap visualization of the 26 shared DEGs revealed a common CRS-responsive transcriptional component across D1 and D2 neurons, while also indicating that most CRS-associated DEGs were not fully shared between the two populations (Figure 2E).

Because several representative downregulated genes in D2 neurons were associated with activity-related transcriptional and signaling responses, we next visualized selected genes at the individual-gene level. *Egr1* was reduced by CRS in both D1 and D2 neurons, indicating that both neuronal populations were transcriptionally engaged by stress (Figure 2F). In contrast, *Nr4a1*, *Nr4a3*, *Rgs2*, and *Junb* showed significant CRS-associated decreases primarily in D2 neurons (Figures 2G–2J). Full mixed-effects model statistics and post hoc comparisons for Figures 2F–2J are provided in Tables S6 and S7. These results suggest that, despite comparable DEG numbers, CRS-associated decreases in several activity-related genes were more evident in D2 neurons.

Together, these findings show that CRS induces transcriptional responses in both D1 and D2 neurons without a simple quantitative dominance of DEGs in either neuronal population. Instead, the CRS response consists of both shared and cell-type-associated components. This observation motivated pathway-level analysis to determine whether comparable DEG numbers are organized into distinct biological programs in D1 and D2 neurons.

### Pathway-level analysis reveals D2-linked convergence of receptor-trafficking and synaptic signaling programs

Because D1 and D2 neurons showed comparable numbers of CRS-responsive genes, we next asked whether these transcriptional responses were organized into distinct pathway-level patterns. Gene set enrichment analysis (GSEA) was performed using genes ranked by CRS-associated expression changes within each neuronal population, and significantly enriched pathways were defined using an FDR q < 0.05 threshold. To summarize the functional organization of CRS-associated transcriptional attenuation, we visualized selected representative negatively enriched Reactome pathways that met the significance threshold and were chosen based on enrichment strength and biological relevance. Negative normalized enrichment scores (NES) indicate enrichment among genes downregulated in CRS relative to control. Complete GSEA results for D1 and D2 neurons are provided in Tables S8 and S9.

In D1 neurons, selected negatively enriched pathways represented multiple functional domains, including HSP90 chaperone cycle for steroid hormone receptors, RHO GTPase-related signaling, recycling pathway of L1, COPI-independent Golgi-to-ER retrograde traffic, and PINK1-PRKN-mediated mitophagy (Figure 3A and Table S8). These pathways suggest that CRS-associated transcriptional attenuation in D1 neurons involves broad intracellular trafficking-, cytoskeletal-, proteostatic-, and stress-related programs. In D2 neurons, selected negatively enriched pathways included receptor-trafficking and synaptic signaling-related programs, including trafficking of AMPA receptors, EPHB-mediated forward signaling, and clathrin-mediated endocytosis (Figure 3B and Table S9). Thus, although CRS-associated pathway-level responses were not restricted to one neuronal population, the selected D2 pathway set showed a more coherent functional organization around receptor-trafficking and synaptic signaling processes.

**Figure 3.**
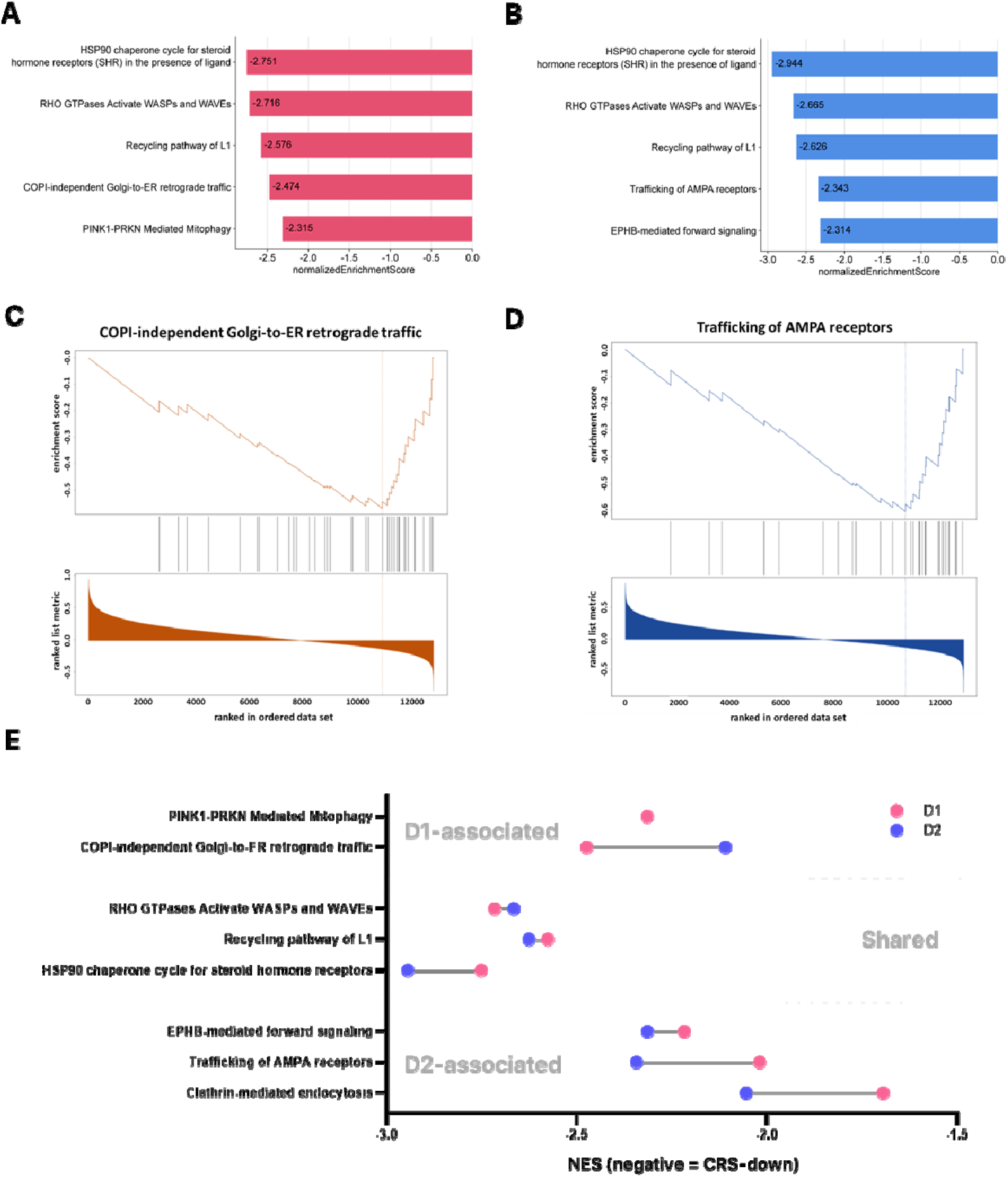
Pathway-level analysis reveals D2-linked convergence of receptor-trafficking and synaptic signaling programs. (A and B) Selected representative negatively enriched Reactome pathways identified by gene set enrichment analysis (GSEA) in D1 neurons (A) and D2 neurons (B). Pathways were selected from significant GSEA results defined by FDR q < 0.05, based on enrichment strength and biological relevance to summarize pathway-level functional organization. Bars indicate normalized enrichment scores (NES). Negative NES values indicate enrichment among genes downregulated in CRS relative to control. (C and D) Representative GSEA enrichment plots showing COPI-independent Golgi-to-ER retrograde traffic in D1 neurons (C) and trafficking of AMPA receptors in D2 neurons (D). These examples were selected from significant negatively enriched pathways to illustrate differences in biological emphasis rather than to indicate strict cell-type specificity. Vertical tick marks indicate the positions of pathway genes within the ranked gene list. The lower panels show the ranked gene list metric used for GSEA. (E) Curated comparison of selected significant negatively enriched pathways grouped by biological theme and relative enrichment profile. Pathway groups summarize D1-linked intracellular/stress-related programs, broadly represented pathway responses, and D2-linked receptor-trafficking/synaptic signaling programs. Grouping was based on functional organization and downstream interpretability rather than strict presence or absence in only one neuronal population. Connected points indicate pathways detected in both neuronal populations, whereas single points indicate pathways detected in only one population under the predefined GSEA threshold of FDR q < 0.05. Pink and blue points indicate NES values in D1 and D2 neurons, respectively. NES, normalized enrichment score; negative NES indicates CRS-downregulated enrichment.

Representative enrichment plots further illustrated selected pathway-level examples. COPI-independent Golgi-to-ER retrograde traffic was shown as a representative intracellular trafficking-related pathway in D1 neurons (Figure 3C), whereas trafficking of AMPA receptors was shown as a representative receptor-trafficking-related pathway in D2 neurons (Figure 3D). These examples were selected to illustrate differences in biological emphasis rather than to indicate strict cell-type specificity. Together, they support the view that CRS engages both neuronal populations, while receptor-trafficking and synaptic signaling programs form a prominent component of the D2 pathway-level response.

To compare the functional organization of selected pathways between neuronal populations, we curated representative significant pathways according to their biological themes and relative enrichment profiles (Figure 3E). D1-linked pathways included PINK1-PRKN-mediated mitophagy and COPI-independent Golgi-to-ER retrograde traffic, consistent with intracellular trafficking-, proteostatic-, and stress-related responses. Shared pathways included RHO GTPase-related signaling, recycling pathway of L1, and HSP90 chaperone cycle, indicating that CRS induced partially shared pathway-level attenuation in both D1 and D2 neurons. In contrast, D2-linked pathways were organized around receptor-trafficking and synaptic signaling processes, including AMPA receptor trafficking, EPHB-mediated signaling, and clathrin-mediated endocytosis.

Together, these results indicate that CRS-associated pathway responses are not completely separated by neuronal population. Rather, GSEA revealed broad pathway-level attenuation in both D1 and D2 neurons, with receptor-trafficking and synaptic signaling programs forming a more coherent D2-linked functional theme. This pathway-level organization provided the rationale for subsequent functional annotation of threshold-defined CRS-responsive genes to determine whether these signals resolved into synapse-centered gene-level categories.

### CRS-downregulated D2 genes are associated with synaptic and postsynaptic functional categories

To determine whether the D2-linked receptor-trafficking and synaptic signaling theme observed by GSEA could be resolved at the threshold-defined DEG level, we next performed DAVID functional annotation using CRS-responsive gene sets defined by the same fold-change and nominal *p* value criteria described above, with upregulated and downregulated genes analyzed separately. Under the same criteria, D1 total and upregulated CRS-responsive gene sets did not show a comparable synapse-centered annotation pattern (Tables S10 and S11), and D1 downregulated genes did not yield significant functional annotation terms. By contrast, D2 CRS-responsive genes yielded functional annotation clusters (Table S12). When D2 genes were separated by direction, upregulated genes were associated with categories such as cilium assembly, chromatin remodeling, cell–matrix adhesion, and potassium ion transmembrane transport (Figure 4A; Table S13), whereas CRS-downregulated D2 genes showed enrichment for synapse-related categories, including glutamatergic synapse and dendritic spine (Figure 4A; Table S14).

**Figure 4.**
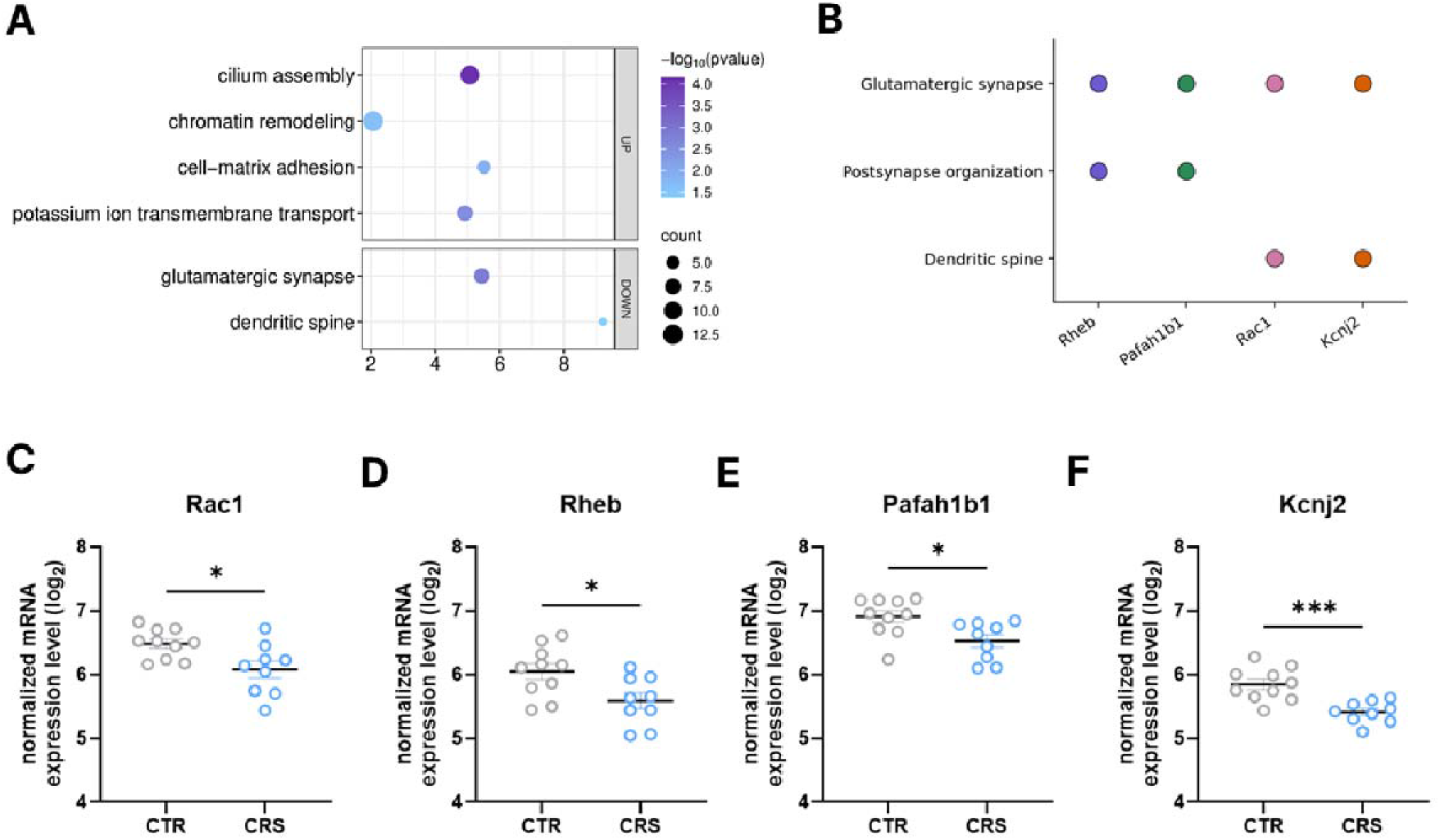
CRS-downregulated D2 genes are associated with synaptic and postsynaptic functional categories. (A) DAVID functional annotation of CRS-responsive genes in D2 neurons. CRS-responsive genes were defined using the same threshold-based DEG criteria as in Figure 2, namely at least a 1.3-fold change and nominal *p <* 0.05. Upregulated and downregulated D2 gene lists were analyzed separately. Selected representative enriched functional categories passing an FDR q < 0.05 threshold are shown. Dot color indicates −log□□(nominal p value), dot size indicates the number of genes associated with each term, and the x-axis indicates enrichment. Full nominal p values and FDR q values are provided in Tables S12–S14. (B) Gene-term association plot showing representative CRS-downregulated D2 genes contributing to selected synapse-related functional terms. Terms were selected from the functional annotation clusters of CRS-downregulated D2 genes to represent nonredundant synaptic domains, including glutamatergic synapse, postsynapse organization, and dendritic spine. Dots indicate gene-term associations. (C–F) Individual gene-level visualization of representative synapse-associated D2 genes. Rac1 (C), Rheb (D), Pafah1b1 (E), and Kcnj2 (F) showed CRS-associated decreases in D2 neurons. Data are shown as log□-normalized expression values and presented as mean ± SEM. Statistical significance was determined by two-tailed unpaired Student’s t test: *Rac1*, *t*(17) = 2.720, *p* = 0.0143; *Rheb*, *t*(17) = 2.563, *p* = 0.0201; *Pafah1b1*, *t*(17) = 2.778, *p* = 0.0129; *Kcnj2*, *t*(17) = 4.277, *p* = 0.0005. \**p* < 0.05, \*\*\**p* < 0.001.

To identify genes contributing to the D2 synapse-related annotation pattern, we examined gene-term associations among representative synapse-related functional terms selected from the functional annotation clusters of CRS-downregulated D2 genes. These terms were chosen to capture nonredundant synaptic domains related to glutamatergic synapse, postsynaptic organization, and dendritic spine structure. This analysis linked *Rheb*, *Pafah1b1*, *Rac1*, and *Kcnj2* to glutamatergic synapse, postsynapse organization, and dendritic spine categories (Figure 4B; Table S14). These genes were selected because they contributed to synapse-related functional terms and showed CRS-associated decreases in D2 neurons.

Individual gene-level visualization further showed that *Rac1*, *Rheb*, *Pafah1b1*, and *Kcnj2* were significantly reduced in D2 neurons following CRS (Figures 4C–4F). These genes are associated with synaptic organization, postsynaptic signaling, dendritic spine regulation, and neuronal excitability-related functions. Thus, although GSEA indicated pathway-level attenuation in both D1 and D2 neurons, DAVID functional annotation of threshold-defined gene sets showed that the D2 downregulated gene set had a clearer synapse-centered functional annotation pattern.

Together, these findings connect the D2-linked receptor-trafficking and synaptic signaling theme observed in Figure 3 to specific synapse-related functional categories and representative downregulated genes. This supports the interpretation that CRS-associated D2 transcriptional attenuation is not defined simply by DEG number, but rather by the organization of downregulated genes around synaptic and postsynaptic molecular programs.

## Discussion

In this study, we used RNAscope-guided GeoMx spatial transcriptomics to compare chronic stress-associated transcriptional responses in D1 and D2 neuronal populations within matched dorsal striatal tissue regions. CRS engaged both neuronal populations and produced comparable numbers of differentially expressed genes in D1 and D2 neurons. However, the biological organization of these responses differed between the two populations. GSEA revealed pathway-level attenuation in both D1 and D2 neurons, but receptor-trafficking and synaptic signaling-related pathways were more clearly supported by synapse-centered functional annotation of CRS-downregulated genes in D2 neurons. These findings suggest that chronic stress does not simply produce a larger transcriptional response in one dorsal striatal cell type. Rather, it reshapes stress-responsive gene programs in a cell-type-associated manner.

The dorsal striatum has historically been studied in relation to action selection, habit formation, and motor control, whereas stress-related affective dysfunction has more often been framed around corticolimbic and ventral striatal circuits^4^. Recent studies, however, increasingly support the relevance of the dorsal striatum to depression- and stress-related behavioral regulation. Human imaging studies have linked depression to altered dorsal corticostriatal and frontostriatal connectivity, striatal dopamine deficits, and impaired reward prediction-error signaling^1–3,5^. Experimental studies further indicate that dorsal striatal circuits contribute to stress coping, reinforcement learning, action–outcome control, habit-related behavioral regulation, and chronic stress-induced circuit remodeling^6–8,13,14^. Together, these studies suggest that the dorsal striatum should not be viewed only as a motor or habit-related structure, but also as a stress-sensitive circuit region that may shape adaptive and maladaptive behavioral responses.

Building on this emerging view of the dorsal striatum as a stress-sensitive circuit region, our findings define a cell-type-associated transcriptional layer of CRS-induced dorsal striatal adaptation. CRS engaged both D1 and D2 neurons, and pathway-level transcriptional attenuation was observed in both populations. The key distinction was organizational: D2 neurons showed a clearer connection between receptor-trafficking and synaptic signaling pathways and synapse-centered functional annotation of threshold-defined CRS-downregulated genes. Thus, in this dataset, the D2-associated stress response was not defined by a larger number of DEGs, but by the functional coherence of downregulated transcriptional programs.

The synapse- and receptor-trafficking-related programs identified here may be relevant to dorsal striatal mechanisms of stress-related behavioral adaptation. AMPA receptor trafficking, EPHB-mediated signaling, glutamatergic synapse, postsynaptic organization, and dendritic spine-related categories point toward molecular processes involved in excitatory synaptic regulation and structural plasticity^15–18^. These results should not be interpreted as direct evidence of altered synaptic transmission, receptor trafficking, or dendritic spine morphology, because these features were not directly measured in the present study. Instead, they provide a spatially resolved transcriptomic basis for future studies testing whether chronic stress alters dorsal striatal synaptic physiology in a cell-type- and circuit-specific manner.

A major next step will be to connect these transcriptional signatures to dorsal striatal circuit function and behavioral outcomes. The dorsal striatum is embedded in cortico-basal ganglia-thalamic loops that support action selection, reinforcement learning, habit formation, and behavioral flexibility^4^. Human imaging and stress-related circuit studies further implicate dorsal striatal interactions with corticostriatal, frontostriatal, amygdala-striatal, and midbrain-striatal pathways in affective and reward-related behavioral regulation^1–3,5,19^. In animal studies, chronic stress has also been shown to alter dorsal striatal microcircuit function, basal ganglia pathway activity, and dorsal striatum–raphe circuit regulation^8,20,21^. Changes in the connectivity, activity, or synaptic efficacy of these circuits could influence stress coping, reinforcement learning, action selection, habit formation, and behavioral flexibility. Therefore, future work should determine how chronic stress alters the functional coupling between dorsal striatal D1/D2 neuronal populations and their upstream or downstream circuit partners. Projection-specific electrophysiology, calcium imaging, circuit tracing, and cell- type-specific gene perturbation will be important for testing whether the transcriptional programs identified here contribute causally to stress-related behavioral adaptation.

In this context, our recent nucleus accumbens study provides complementary evidence that CRS-associated D2R-expressing neuronal molecular states can be linked to depressive-like behavioral outcomes^22^. However, that study was not designed to directly compare D1 and D2 transcriptional responses within matched striatal tissue regions. The present study extends this line of investigation to a different striatal territory by directly comparing D1 and D2 neuronal transcriptional organization within matched dorsal striatal ROIs. Together, these studies support a broader model in which chronic stress reshapes striatal molecular states in a region- and cell-type-associated manner. This model does not imply that one striatal cell type or region alone accounts for stress-related behavioral dysfunction. Instead, it highlights the need for integrative studies that connect molecular organization, circuit activity, and behavioral output across ventral and dorsal striatal territories.

Thus, the major contribution of this study is not simply the identification of CRS-responsive genes in the dorsal striatum, but the spatially matched comparison of how these genes are organized across D1 and D2 neuronal compartments. By profiling both populations within matched tissue regions, our analysis highlights cell-type-associated differences in the functional organization of stress-responsive transcriptional programs that would not be captured by DEG number alone. This within-region framework may be useful for future studies seeking to connect stress-associated transcriptional states to synaptic physiology, circuit function, and behavioral adaptation in anatomically defined striatal territories.

Several limitations should be considered. First, RNAscope-guided GeoMx profiling provides transcriptomic information from D1- and D2-enriched molecular compartments within selected dorsal striatal ROIs, but it does not resolve transcriptomes at single-cell resolution. Second, the present study was designed to define spatially matched D1/D2 transcriptional organization in the dorsal striatum, and the functional consequences of the identified transcriptional programs remain to be tested directly. Third, the analysis was performed in male mice under a CRS paradigm and within a defined anterior dorsal striatal sampling range; therefore, future studies should examine whether similar or distinct D1/D2 transcriptional organization occurs across sex, dorsal striatal subregions, developmental stress exposure, stress models, and behavioral time points^23,24^. Despite these limitations, our findings provide a framework for future studies aimed at linking dorsal striatal cell-type-associated transcriptional programs to synaptic physiology, circuit function, and stress-related behavioral regulation.

## Resource availability

### Lead contact

Further information and requests for resources and reagents should be directed to and will be fulfilled by the lead contact, Heh-In Im.

### Materials availability

This study did not generate new unique reagents.

### Data and code availability

Raw and processed GeoMx spatial transcriptomic data generated in this study, including FASTQ files, DCC files, the associated PKC file, and the Q3-normalized expression matrix, have been deposited in the NCBI Gene Expression Omnibus (GEO) and are publicly available under accession number GSE337812. For convenience, the Q3-normalized expression matrix, DEG lists, pathway enrichment results, and functional annotation results are also provided in the supplemental tables.

This paper does not report original code.

Any additional information required to reanalyze the data reported in this paper is available from the lead contact upon reasonable request.

## Acknowledgments

This work was supported by the Ministry of Food and Drug Safety (grant no. 25212MFDS003), the National Research Foundation of Korea (grant nos. RS-2024-00332024 and RS-2024-00463082), and the Korea Institute of Science and Technology (grant no. 26Z9001). We thank Eonji Noh and Seungyoun Lee of Theragen Bio for their assistance with bioinformatics analysis. Brain diagrams were adapted from The Mouse Brain in Stereotaxic Coordinates (Paxinos and Franklin) and the Allen Brain Atlas (https://atlas.brain-map.org/). Some figures were created using BioRender.com.

## Author contributions

J.B. and H.-I.I. conceived and designed the study. J.B. performed the experiments, analyzed the data, generated the figures, and wrote the initial draft of the manuscript. H.-I.I. supervised the study, provided conceptual guidance, secured funding, and edited the manuscript. Both authors reviewed and approved the final manuscript.

## Supplemental information

Supplemental information includes Figures S1–S5 and Tables S1–S14. Supplemental figures are provided in the supplemental information file, and supplemental tables are provided as an Excel workbook.

## Methods

### Animals

Male heterozygous Drd2-Cre mice, B6.FVB(Cg)-Tg(Drd2-cre)ER44Gsat/Mmcd (#032108-UCD; MMRRC, CA, USA), were used at 11–13 weeks of age, consistent with related CRS behavioral experiments performed in this mouse background. Mice were singly housed throughout the experimental period under a 12-h reversed light/dark cycle, with lights off at 8:00 AM and lights on at 8:00 PM. Food and water were available ad libitum. Cre activity was not used for labeling or genetic manipulation in this study; D1 and D2 neuronal segments were defined independently based on endogenous *Drd1* and *Drd2* RNAscope signals. All animal procedures were approved by the Institutional Animal Care and Use Committee of the Korea Institute of Science and Technology (KIST-IACUC-2024-072-9).

### Chronic restraint stress procedure

Chronic restraint stress was performed using a rectangular restraint device with an internal dimension of 3 × 3 × 17 cm. Mice were positioned in the device, and a movable fixation wall was adjusted to approximately 4 cm to restrict forward and backward movement while allowing limited postural adjustment. CRS mice were restrained simultaneously for 6 h per day for 21 consecutive days. Control mice were not exposed to restraint stress and were handled for approximately 3 min per day during the same period. Cage maintenance and body weight measurements were conducted regularly for both groups throughout the CRS paradigm. Brains were collected 3 days after the final restraint session, matching the post-CRS time point at which behavioral phenotypes had been validated in prior experiments.

### Formalin-fixed paraffin-embedded tissue preparation

Mouse brains were harvested and immersed in 4% paraformaldehyde (PFA) at 4°C for 24 h. After fixation, tissues were dehydrated sequentially in 40% ethanol, 70% ethanol, and fresh 70% ethanol for 24 h at each step. Fixed brains were placed in embedding cassettes and processed using a Tissue-Tek VIP Tissue Processor (Sakura Finetek). The tissue-processing program consisted of a 30-min ethanol wash, five 1-h ethanol incubations, three 45-min CitriSolv washes (Decon Labs), and three 1-h paraffin infiltration steps over a 12-h cycle.

Following paraffin embedding, 2-mm-diameter tissue cores containing the dorsal striatum were obtained from individual paraffin blocks and re-embedded into tissue microarray (TMA) blocks. This TMA-based preparation enabled simultaneous processing of multiple samples under identical staining, hybridization, and GeoMx DSP profiling conditions. Paraffin-embedded tissues were sectioned at 5 μm thickness for RNAscope staining and GeoMx DSP analysis.

### RNAscope-guided D1/D2 segmentation and GeoMx spatial transcriptomic profiling

RNAscope-based fluorescence labeling was performed on formalin-fixed paraffin-embedded brain sections using the RNAscope Multiplex Fluorescent Reagent Kit v2 (Advanced Cell Diagnostics, ACD) according to the manufacturer’s instructions. Briefly, tissue sections were baked at 60°C for 1 h, deparaffinized, and subjected to sequential pretreatment steps, including hydrogen peroxide treatment, target retrieval, and protease digestion. Sections were then incubated with probes targeting *Drd1* (ACD, 461901) and *Drd2* (ACD, 406501-C2) at 40°C for 2 h. Probe signals were amplified using the recommended AMP reagents and detected with Opal 520 and Opal 620 fluorophores (Akoya Biosciences). After RNAscope labeling, sections were post-fixed and processed for GeoMx Digital Spatial Profiler analysis following NanoString’s standard workflow.

For spatial transcriptomic profiling, sections were hybridized overnight at 37°C with the GeoMx Mouse Whole Transcriptome Atlas probe set (NanoString). After hybridization, slides were washed, incubated with Buffer W for 30 min, and stained with a visualization cocktail containing SYTO 83 nuclear dye for 2 h at room temperature in a humidified chamber.

Fluorescence images were acquired using the GeoMx DSP platform to guide region-of-interest selection. Dorsal striatal ROIs were manually selected based on tissue morphology, nuclear staining, RNAscope probe signals, and bregma-referenced anatomical landmarks. ROIs were selected from sagittal sections at approximately ML 1.0–1.3 mm lateral to the midline, within the dorsal striatum corresponding approximately to AP +1.1 to +1.4 mm from bregma and DV 2.0–3.0 mm. Each mouse contributed one dorsal striatal ROI, and each ROI was segmented into two molecular compartments corresponding to D1 and D2 neuronal populations. Thus, each mouse provided one ROI and paired D1 and D2 segments for within-region comparison. ROIs were included only when both D1 and D2 compartments contained at least 200 nuclei. This ROI-level inclusion criterion was applied in addition to the standard segment-level GeoMx QC threshold described below. Within each ROI, D1 and D2 neuronal compartments were defined using endogenous *Drd1* and *Drd2* transcript signals, respectively. D1/D2 segmentation was performed independently of Cre activity.

Photocleavable indexing oligonucleotides from each segmented compartment were released by UV illumination and collected into a 96-well plate by microcapillary aspiration. The collected oligonucleotides were used for library preparation and sequenced on an Illumina NovaSeq 6000 platform by Theragen Bio (Seongnam, Korea).

### Sequencing data processing and quality control

Sequencing reads from GeoMx DSP libraries were processed using the GeoMx NGS analysis pipeline as previously described. Briefly, FASTQ files were quality checked, adapter trimmed, merged into stitched reads, aligned to probe-specific barcodes, and deduplicated using unique molecular identifiers. Deduplicated counts were imported into the GeoMx DSP Data Analysis Suite for quality control and downstream analysis.

Segment QC required ≥1,000 raw reads, ≥50% alignment rate, ≥80% stitched and trimmed reads, ≥50% sequencing saturation, geometric mean of negative probe counts ≥1, no-template control counts <2,000, tissue area ≥1,000 μm², and ≥50 nuclei per segment. The ≥50-nuclei threshold represents the standard segment-level GeoMx QC criterion, whereas ROI inclusion required at least 200 nuclei in each D1 and D2 segment before collection. Underperforming probes were removed when their mean count was ≤10% of the total probe counts for the corresponding gene, and genes were excluded when expression was below the limit of quantitation in more than 20% of all D1 and D2 segments included in the final dataset. The LOQ was calculated as GeoMean (Negative probes) × [GeoSD (Negative probes)]². Filtered count data were normalized using Q3 normalization.

### Transcriptomic and pathway analyses

Transcriptomic analyses were performed using samples from 10 control mice and 9 CRS mice. Each mouse contributed one dorsal striatal ROI, which was segmented into one D1 and one D2 neuronal compartment. Thus, each mouse provided paired D1 and D2 segment-level expression profiles, and the mouse, rather than the segment, was treated as the biological replicate for gene-level statistical analyses.

Gene-level expression analyses were performed using log□J-transformed Q3-normalized expression values. For each gene, log□J fold change was calculated as the difference between the mean log□J-normalized expression value in the CRS group and that in the control group within each neuronal population:

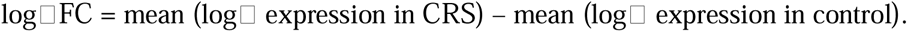

Fold change values, when shown, were derived as 2^(log□FC)^. For threshold-based differential expression analyses, upregulated genes were defined as genes with log□JFC ≥ 0.3785 and nominal *p* < 0.05, corresponding to FC ≥ 1.3. Downregulated genes were defined as genes with log□JFC ≤ −0.3785 and nominal *p* < 0.05, corresponding to FC ≤ 0.77. For more stringent filtering, genes with |log□JFC| ≥ 1 and nominal *p* < 0.05 were used where indicated, corresponding to at least a twofold change. These threshold-defined CRS-responsive gene sets were used for DEG summary, overlap analysis, heatmap visualization, and downstream functional annotation.

Gene set enrichment analysis (GSEA) was performed separately from threshold-defined DEG analysis using ranked gene lists without applying an FC cutoff. GSEA was performed using WebGestalt 2024 (https://www.webgestalt.org/) with Reactome gene sets. Genes were ranked by log□JFC values comparing CRS with control conditions within each neuronal population. Gene symbols were used as input identifiers, and duplicate gene symbols were collapsed using the mean value. Gene sets containing 20–2,000 genes were included in the analysis. Enrichment was assessed using 1,000 permutations, and pathways with FDR q < 0.05 were considered significant. Redundancy among enriched categories was reduced using the weighted set cover method. Normalized enrichment score (NES) was used to represent the direction and magnitude of enrichment. Negative NES values indicate enrichment among genes downregulated in CRS relative to control.

Gene ontology enrichment analysis was performed using ToppFun in the ToppGene Suite (https://toppgene.cchmc.org/). Threshold-defined gene lists were analyzed to identify enriched GO categories, and terms with FDR q < 0.05 were considered significant. For visualization, enriched terms passing the FDR threshold were displayed using −log□□ (nominal p value). Full nominal p values and FDR q values are provided in the supplemental tables.

Functional annotation of selected gene sets was additionally performed using DAVID Bioinformatics Resources (https://david.ncifcrf.gov/). DAVID functional annotation results were evaluated using FDR-adjusted values where applicable, and categories with FDR q < 0.05 were considered significant. For the analyses presented in Figure 4, D1 and D2 CRS-responsive gene lists were analyzed using the same DEG and enrichment criteria, with upregulated and downregulated gene lists analyzed separately where indicated. Synapse- related categories emerging from CRS-downregulated D2 genes were further examined at the gene-term level.

Protein-protein interaction (PPI) network analysis was performed and visualized using STRING (https://string-db.org/) with Mus musculus as the reference organism. Selected genes associated with representative GO or functional annotation categories were submitted to STRING to visualize gene-level organization within enriched categories. Networks were generated using STRING functional association networks with the combined interaction score, and interactions were displayed using a minimum required interaction score of 0.400, corresponding to medium confidence. For baseline D1–D2 characterization under control conditions, genes associated with representative synapse-related categories were analyzed to compare network composition between D1 and D2 neurons.

### Statistical analysis

Statistical analyses of individual gene expression values were performed using mouse-level log□-normalized expression values. For comparisons between control and CRS groups within a single neuronal population, two-tailed unpaired Student’s t tests were used. For paired comparisons between D1 and D2 segments derived from the same ROI, paired two-tailed Student’s t tests were used where appropriate. For analyses comparing the effects of stress condition and neuronal population simultaneously, stress condition was treated as a between-subject factor and neuronal population or segment identity as a within-subject factor. Two-way ANOVA or mixed-effects models were used with stress condition and neuronal population as factors, and mouse identity was used to account for within-mouse pairing in analyses involving paired D1/D2 segments, followed by appropriate post hoc comparisons where indicated. Statistical analyses and graph generation were performed using GraphPad Prism 11. Data are presented as mean ± SEM unless otherwise stated. A nominal p value < 0.05 was considered statistically significant for individual gene-level comparisons.

