## Supplementary material for "Spatial transcriptomics reveals D2-associated synaptic transcriptional attenuation in the chronically stressed dorsal striatum": supplemental information.pdf

Jinhee Bae[1] and Heh-In Im[1,2]*

[1]Center for Brain Disorders, Brain Science Institute, Korea Institute of Science and Technology, Seoul, South Korea, 02792.

[2]Division of Bio-Medical Science & Technology, KIST School, Korea University of Science and Technology, Seoul, South Korea, 02792.

This file includes Figures S1–S5 and supplemental figure legends.

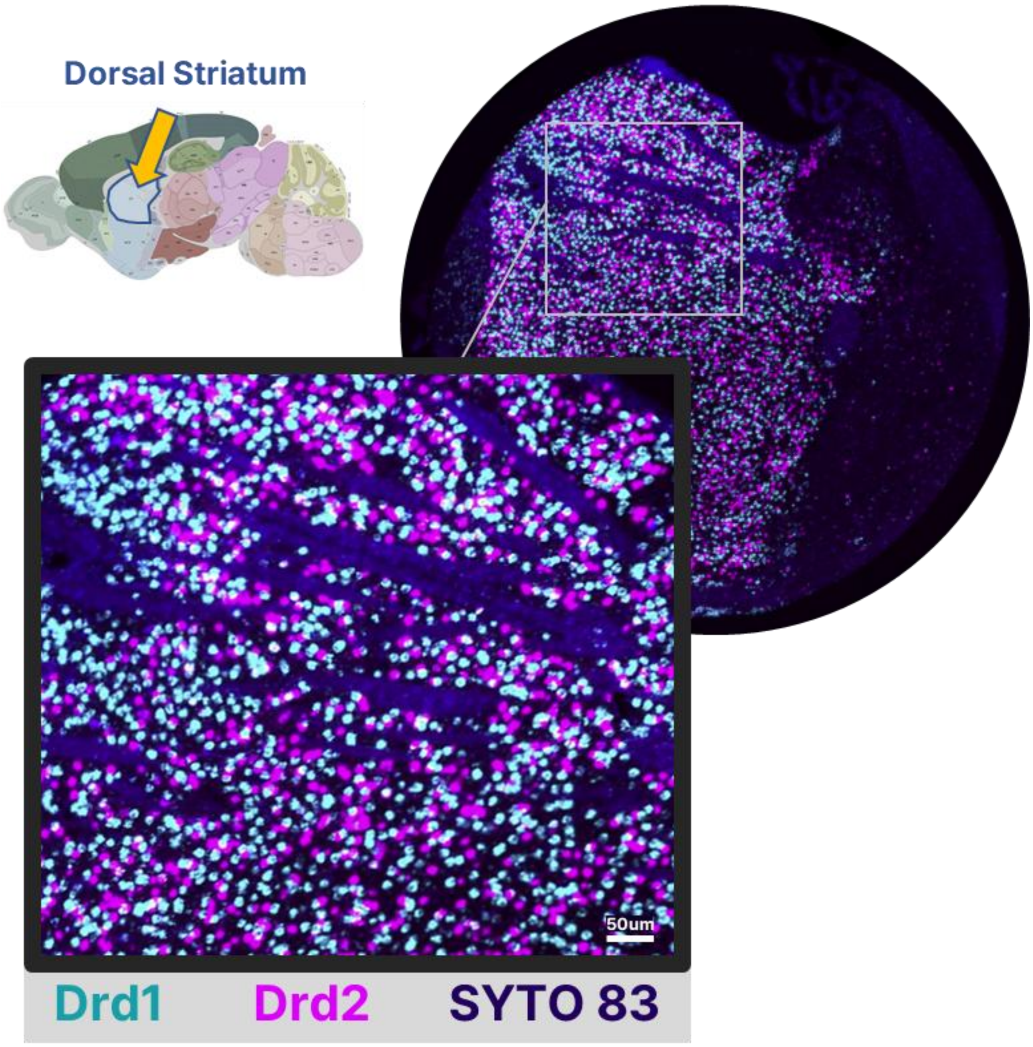


**Figure S1. Validation of RNAscope-guided D1 and D2 neuronal segmentation in the dorsal striatum.**

Representative RNAscope image showing *Drd1*-, *Drd2*-, and SYTO 83-labeled cells in the dorsal striatum. The magnified image shows clearly distinguishable *Drd1* and *Drd2* signals used for D1/D2-informed segmentation in GeoMx spatial transcriptomic profiling. Scale bar, 50 µm.

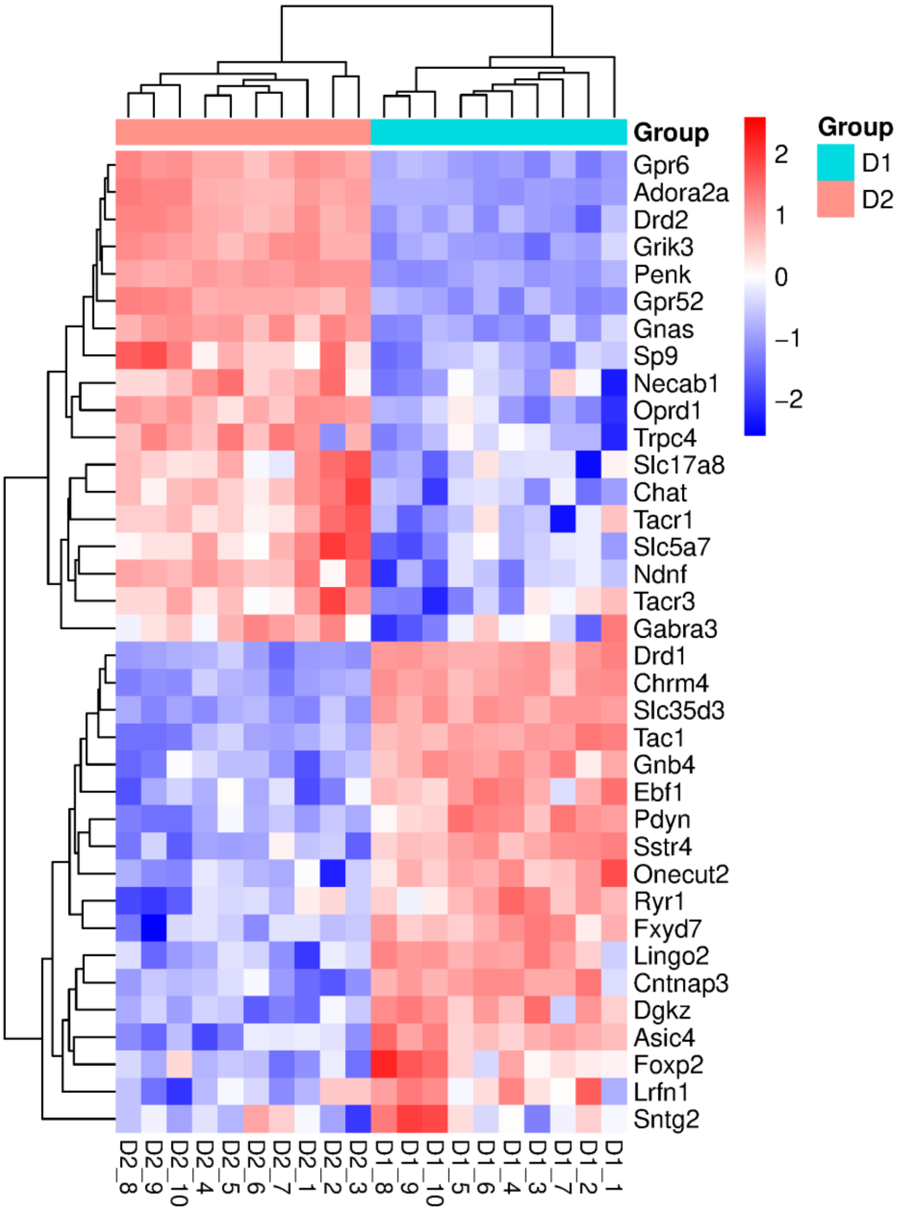


**Figure S2. Baseline marker-based characterization of D1 and D2 neuronal segments.**

Heatmap showing expression patterns of representative D1- and D2-associated genes under control conditions. D1-associated genes, including *Drd1*, *Pdyn*, *Tac1*, and *Chrm4*, and D2-associated genes, including *Drd2*, *Penk*, *Adora2a*, and *Gpr6*, showed expected enrichment patterns in the corresponding segmented populations. Rows represent genes, and columns represent individual D1 or D2 segments. Color indicates row-scaled normalized expression.

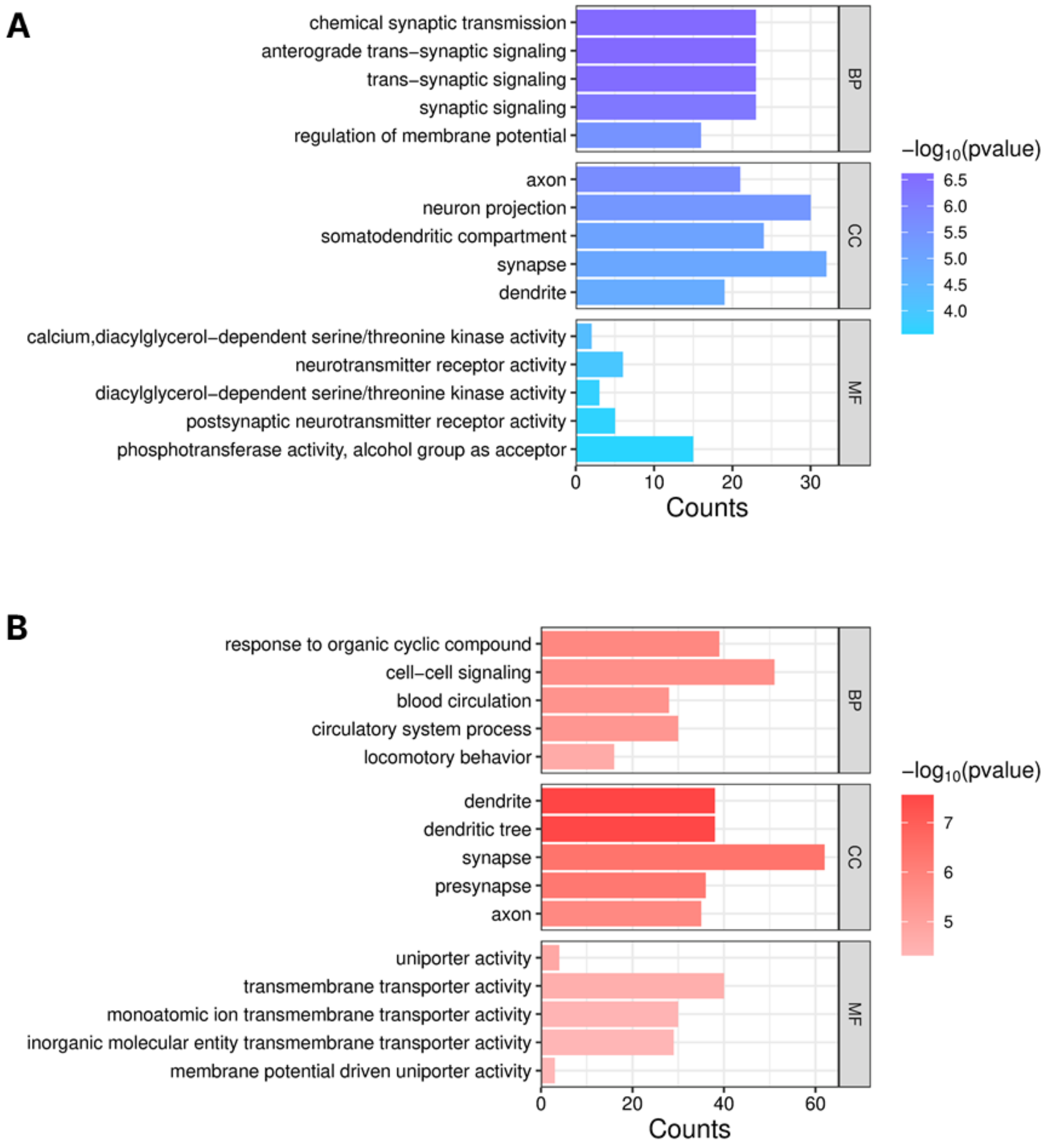


**Figure S3. Baseline functional annotation of shared GO categories in D1 and D2 neuronal segments.**

(A and B) Gene ontology enrichment analysis of representative baseline D1- and D2-associated gene sets under control conditions. GO terms identified in both D1 and D2 analyses are plotted separately to compare functional annotation patterns between D1-associated genes (A) and D2-associated genes (B). Enriched GO terms were defined using an FDR $q < 0.05$ threshold. Terms are grouped by biological process (BP), cellular component (CC), and molecular function (MF). Bar length indicates gene counts, and color indicates

$-\log_{10}$(nominal $p$ value). Full nominal $p$ values and FDR $q$ values are provided in Tables S2 and S3. These analyses show that D1 and D2 segments may share broad neuronal and synapse-related GO labels while being supported by distinct gene sets.

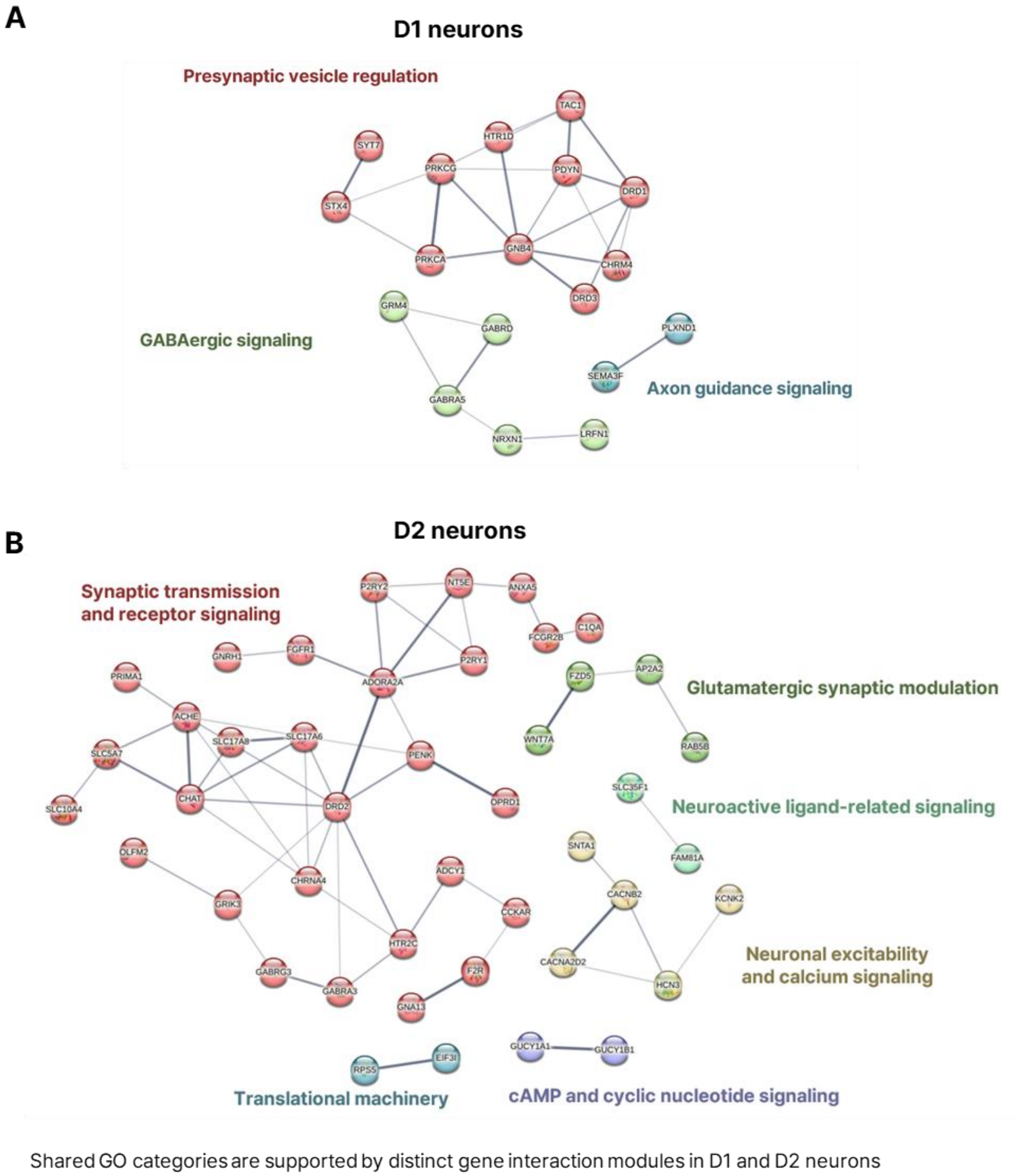


**Figure S4. PPI networks of genes associated with shared synapse-related GO terms in D1 and D2 neuronal segments under control conditions.**

STRING protein-protein interaction (PPI) networks showing genes associated with representative shared synapse-related GO categories in D1- and D2-associated baseline gene sets. Networks were generated separately for D1-associated and D2-associated gene sets to compare gene composition within broadly overlapping synapse-related functional labels.

STRING analysis was performed using *Mus musculus* as the reference organism, and interactions were visualized using STRING functional association networks with the combined interaction score at medium confidence or higher (score ≥ 0.400). Functional modules were annotated based on gene composition and biological relevance. These analyses show that shared GO terms can be supported by distinct gene-level network structures in D1 and D2 neuronal populations.

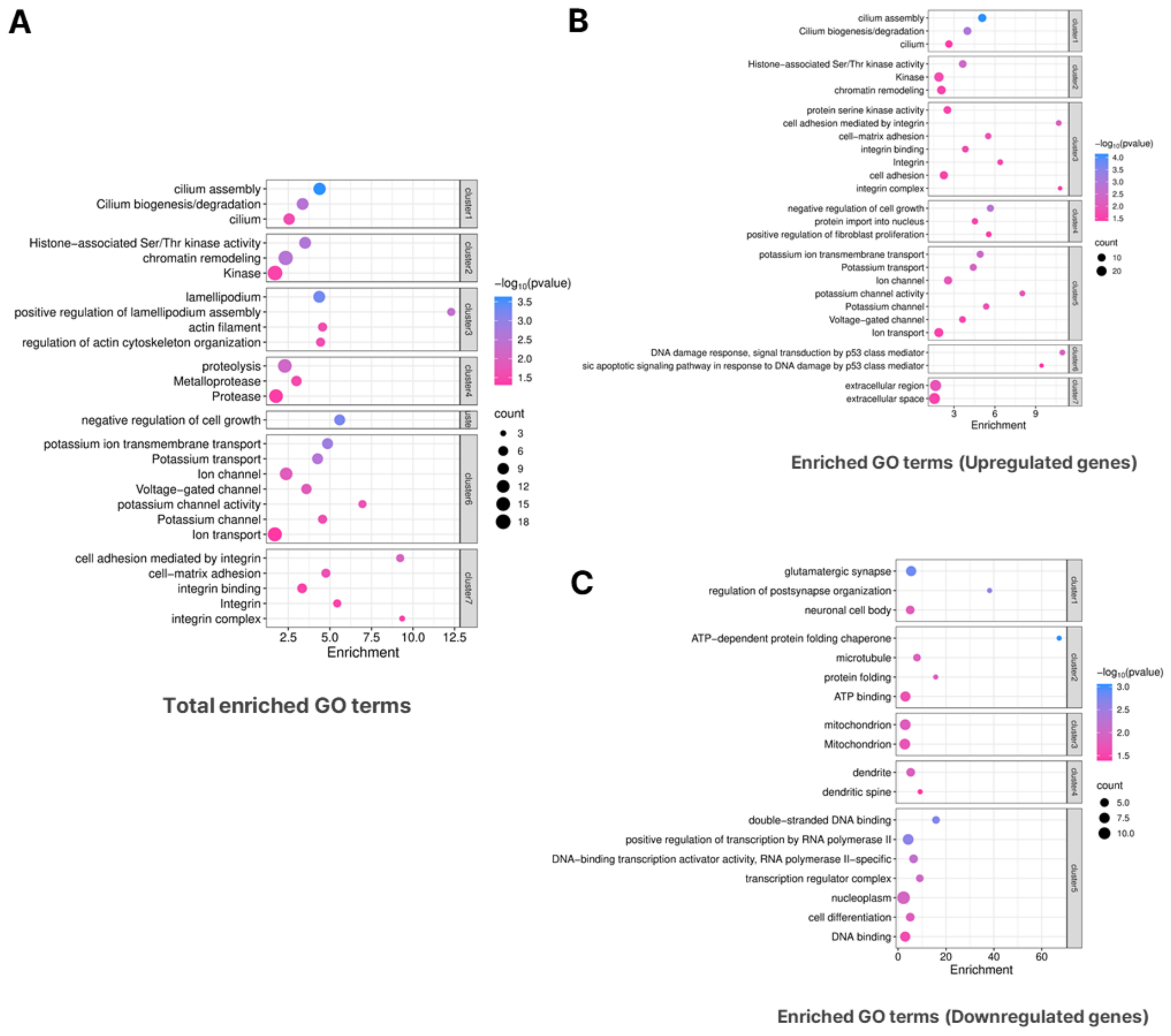


**Figure S5. Extended functional annotation of CRS-responsive genes in D2 neurons.**

(A) DAVID functional annotation of all CRS-responsive genes in D2 neurons. (B and C) DAVID functional annotation of upregulated (B) and downregulated (C) CRS-responsive genes in D2 neurons. CRS-responsive genes were defined using the same threshold-based DEG criteria as in Figure 2, namely at least a 1.3-fold change and nominal $p < 0.05$. Representative enriched functional categories passing an FDR $q < 0.05$ threshold are shown. Dot color indicates $-\log_{10}$(nominal $p$ value), dot size indicates the number of genes associated with each term, and the x-axis indicates enrichment. Full nominal $p$ values and FDR $q$ values are provided in Tables S12–S14. Downregulated D2 genes included synapse-related categories such as glutamatergic synapse and dendritic spine, consistent with the D2-associated synaptic transcriptional attenuation observed in the main analyses.
